# ACE2-independent interaction of SARS-CoV-2 spike protein to human epithelial cells can be inhibited by unfractionated heparin

**DOI:** 10.1101/2020.05.21.107870

**Authors:** Lynda J. Partridge, Lucy Urwin, Martin J.H. Nicklin, David C. James, Luke R. Green, Peter N. Monk

**Author notes:** Corresponding author (LRG).

## Abstract

The SARS-CoV-2 spike protein is known to bind to the receptor, ACE2, on the surface of target cells. The spike protein is processed by membrane proteases, including TMPRSS2, and either internalises or fuses directly with the cell, leading to infection. We have identified a human cell line that expresses both ACE2 and TMPRSS2, the RT4 urinary bladder transitional carcinoma, and used it to develop a proxy assay for viral interactions with host cells. A tagged recombinant form of the spike protein, containing both the S1 and S2 domains, interacted strongly with RT4 cells as determined by flow cytometry, whereas the S1 domain and the receptor binding domain (RBD) interacted weakly. S1S2 interaction was temperature dependent and increased sharply at 37°C, suggesting that processing of the intact spike protein is likely to be important in the interaction. S1S2 protein could associate with cells with a low dependence on ACE2 expression, while RBD required the presence of ACE2 for interaction. As the spike protein has previously been shown to bind heparin, a soluble glycosaminoglycan, we used a flow cytometric assay to determine the effect of heparin on spike protein interaction with RT4 cells. Unfractionated heparin inhibited spike protein interaction with an IC_50_ value of <0.05U/ml whereas two low molecular weight heparins were much less effective. A mutant form of the spike protein, lacking the Arg-rich region proposed to be a furin cleavage site, interacted very weakly with cells and had a lower affinity for unfractionated and lower molecular weight heparin than the wild type spike protein. This indicates that the furin cleavage site might also be a heparin binding site and potentially important in interactions with host cells. Taken together, our data suggest that heparin, particularly unfractionated forms, could be considered to reduce clinical manifestations of COVID-19 by inhibiting continuing viral infection.

**Author Summary:** Since the emergence of SARS-CoV-2 in 2019, the world has faced a vast public health crisis. SARS-CoV-2 associates with human cells through interaction of the viral spike protein with the host receptor, ACE2. In the absence of a vaccine, new treatments are required to reduce the morbidity and mortality of SARS-CoV-2. Here, we use a novel technique to demonstrate spike protein interactions with human cells with low levels of ACE2 at the cell surface, suggesting a secondary receptor. We demonstrate the importance of a new heparin-binding site within the viral spike protein for these interactions. We also found that unfractionated heparin was able to bind to the viral spike protein and therefore, potently inhibit viral spike protein interactions with human cells. Our data demonstrate that ACE2 is not absolutely required for spike protein interactions with human cells and furthermore, that unfractionated heparin should be considered as a treatment to reduce SARS-CoV-2 viral infection.

## Introduction

SARS-CoV-2, the causative agent of COVID-19, is thought to infect cells after binding with high affinity to a host cell receptor, ACE2 [1]. The ACE2 binding domain is located in the spike protein that consists of two domains: S1, which has a high affinity receptor binding domain (RBD) and S2, containing sequences necessary for fusion with the host cell. S1 and S2 are linked by a sequence that contains a putative furin cleavage site that is critical for the entry of the virus into human cells [2]. A cell-surface host serine protease, TMPRSS2, is also thought to be involved in viral entry and is proposed to cleave S1 and S2, leading to activation of the fusion machinery [1]. By analogy with SARS-CoV, it is expected that the virus can fuse at the cell surface or later, following internalisation (reviewed in [3]).

Paradoxically, ACE2 is expressed at quite low levels by most cell types [4] and by very few cell lines, leading to suggestions that additional receptor sites must exist. Viruses, such as herpes simplex and the β coronavirus family, are known to interact with host glycosaminoglycans [5]. A growing body of evidence suggests that SARS-CoV-2 can bind the glycosaminoglycans, heparan sulphate and heparin, dependent on their level of sulphation [6–9] and that heparin can inhibit SARS CoV 2 entry into host cells. Initial binding to heparan sulphates is thought to keep the spike protein within an ‘open’ conformation allowing for downstream binding and processing by ACE2 and TMPRSS2 respectively [8].

Here we present a new assay for viral attachment to host cells, using a human bladder epithelial cell line that expresses both ACE2 and TMPRSS2. The intact viral spike protein, but not the isolated S1 or RBD, exhibits a temperature dependent interaction that allows rapid detection by flow cytometry. We have used this assay to confirm that heparin can inhibit spike protein interactions and demonstrate that high affinity heparin binding requires the putative furin cleavage site and the S2 domain of the viral spike protein but is independent of the S1 domain.

## Materials and Methods

### Materials

Unfractionated heparin (Leo, 1000U/ml), dalteparin (25000IU/ml) and enoxaparin (10000IU/ml) were obtained from the Royal Hallamshire Hospital Pharmacy, Sheffield, UK. Fondaparinux was purchased from Merck, UK. Goat anti-human ACE2 antibody AF933 (Biotechne), goat control IgG AB-108-C (Biotechne), rabbit anti-human TMPRSS2 (MBS9215011, Gentaur), rabbit IgG control (Biolegend) were used as per manufacturers’ instructions.

### Cell culture

The RT4 (ATCC® HTB-2™) cell line was obtained from American Tissue Type Collection (ATCC) and cultured in McCoys 5A medium (Thermo Fisher Scientific) supplemented with 10% foetal calf serum (FCS). The A549 cell line was obtained from the European Collection of Animal Cell Cultures (ECACC) and cultured in DMEM supplemented with 10% FCS. The 293T (ATCC® CRL-3216™) cell line was obtained from ATCC and cultured in DMEM supplemented with 10% FCS. 293T_ACE2_ cells were kindly provided by Paul Bieniasz (The Rockefeller University, USA), cultured as described for 293T cells including selection with 5ug/ml blasticidin. A human keratinocyte cell line, HaCaT (300493), was obtained from Cell Line Services (CLS GmbH, Germany) and routinely cultured in DMEM supplemented with 10% FCS. The Caco2 cell line (ATCC® HTB-37), derived from a colorectal adenocarcinoma, was obtained from Dr. Michael Trikic (University of Sheffield, UK). Cell lines were routinely sub-cultured by trypsinisation and maintained in sub-confluent cultures.

### Real time quantitative PCR (RT-qPCR)

HaCaT, RT4, A549, 293T, 293T_ACE2_ and Caco2 cell lines were cultured for 48hrs under standard media conditions and harvested using trypsin. Total RNA was extracted using the RNeasy Mini Kit (Qiagen) and quantified using a NanoPhotometer N60 Touch (Implen). RNA samples were converted to cDNA using the High-Capacity cDNA Reverse Transcription Kit (Applied Biosystems). Control samples containing no reverse transcriptase or no RNA template were included. RT-qPCR was performed by SYBR Green assay using the PrecisionPLUS OneStep RT-qPCR Master Mix and the QuantStudio 5 Real-Time PCR system. Gene expression levels of ACE2 and TMPRSS2 were investigated by Comparative CT experiments and GAPDH expression was measured as an endogenous control. All primers were designed using PrimerBLAST (see Supplementary Table 1 for primer sequences). The A549 cell line has been used as a reference for ACE2/TMPRSS2 gene expression and the data is presented as mean Relative Quantification (RQ).

**Table 1.** Normalised mRNA expression values for ACE2 and potentially associated membrane proteins in several cell lines.

|  | **RT4** | **A549** | **Caco2** | **HaCaT** |
| --- | --- | --- | --- | --- |
| **ACE2** | 2.40 | 0.00 | 0.00 | 4.10 |
| **TMPRSS2** | 35.2 | 0.00 | 14.6 | 0.00 |
| **ADAM17** | 12.9 | 12.7 | 11.2 | 14.2 |
| **RHBDL2** | 6.00 | 18.8 | 5.50 | 13.9 |
| **CD9** | 94.7 | 14.9 | 10.9 | 74.5 |

Normalised mRNA expression values are provided from the Protein Atlas database (https://www.proteinatlas.org).

### Detection of cell surface ACE2 and TMPRSS2 expression

A549 and RT4 cells were harvested using trypsin/EDTA and resuspended in BBN (Hank’s balanced salt solution (HBSS), 0.1% (w/v) Bovine Serum Albumin (BSA), 0.2% (w/v) sodium azide). Cells were incubated with anti-human ACE2 and a goat immunoglobulin control at 15ug/ml, or anti-human TMPRSS2 and a rabbit immunoglobulin negative control at 40ug/ml for 60 min at 37C, then washed twice in BBN. Cell associated immunoglobulin was detected with an appropriate FITC-labelled secondary antibody and fluorescence measured using an Attune NXT cytometer (Thermofisher, UK). Data are shown as relative fluorescence intensities calculated as a percentage of the negative control fluorescence.

### Purification of modified S1S2 and RBD

Spike protein attachment to cells was tested using four versions of the SARS-CoV-2 spike protein. Wild-type S1S2 (wt S1S2; Val16-Pro1213; Stratech UK) with a His6 tag at the C-terminus was expressed in baculovirus-insect cells, while S1-Fc (Val16-Arg685; Stratech UK) with a mouse IgG1 Fc region at the C-terminus was expressed in HEK293 cells. Modified S1S2 (mS1S2) and the receptor binding domain (RBD) cloned into a pCAGGS expression vector were kindly provided by Florian Krammer (Mount Sinai, USA) [10]. A polybasic cleavage site, recognised by furin, was removed (RRAR to A) in mS1S2 and two stabilising mutations were added (K986P and V987P). A thrombin cleavage site, a T4 foldon sequence allowing trimerization and a His6 tag were fused to the C-terminal amino acid P1213. RBD was expressed using the natural signal peptide fused to RBD (aa 419-541) and a His6 tag at the C terminus. Recombinant proteins were expressed and purified as described previously [10].

### Spike protein interaction assay

Cells were harvested by brief trypsinisation and added to wells of a 96-well U bottom plate. After centrifugation at 300xg for 3 min and washing with HBSS containing divalent cations and 0.1% BSA (assay buffer, AB), cells were incubated with potential inhibitors in 50µl AB for 30 min at 37°C. The supernatant was removed following centrifugation and 25µl of AB containing S1-Fc (Stratech UK), RBD-His6 (produced in-house) or S1S2-His6 protein (Stratech UK) added before incubation at 4, 21 or 37°C for 60 min. Cells were washed once and then incubated with the appropriate fluorescently labelled secondary antibody (anti-mouse polyvalent Ig-FITC (Merck) or anti-His6 HIS.H8 DyLight 488, (Invitrogen) for 30 min at 21°C. Cells were finally resuspended in 50µl AB containing propidium iodide and cell-associated fluorescence measured using a Guava 2L-6HT flow cytometer. Live cells were gated as a propidium iodide negative population and the median fluorescence (MFI) recorded. MFI was calculated after subtraction of cell-associated fluorescence of the secondary antibody alone. Where stated, the data were normalised to the untreated control cells.

### Determination of spike protein binding to heparin by ELISA

Heparin-binding plates (Plasso EpranEx™), a gift from Dr David Buttle (University of Sheffield, UK), were coated with 10µg/ml of UFH or dalteparin diluted in phosphate buffered saline (PBS) (150μl/well) at room temperature overnight. Wells were washed twice with PBS 0.05% Tween (PBST), blocked with PBST with 1% BSA for 2 hr at 37 °C before three further washes with PBST. Various concentration of His-tagged spike proteins in AB buffer (or AB buffer control) were added to the wells (50μl/well) and incubated at 37 °C for 2 hr. Wells were washed three times as above then incubated at room temperature with 50μl/well biotin-labelled rabbit monoclonal anti-His6 (Thermo Fisher Scientific) diluted to 1/1000 in AB for 1 hr, washed 3 times with PBST and incubated for 30 min with 50μl/well streptavidin-HRP (Pierce) diluted 1/200 in AB. After washing 3 times with PBST and twice with dH_2_O, 50μl per well TMB substrate solution (Novex) was added followed by 50μl 1M HCl to quench the reaction. Absorbance was measured at OD_450nm_ with non-specific spike protein binding to uncoated wells subtracted.

## Results

### Selection of a cell line for viral spike protein attachment studies

mRNA expression data provided by The Protein Atlas database in a variety of human cell lines was confirmed by quantitative PCR (https://www.proteinatlas.org; [11]). The human lung adenocarcinoma alveolar basal epithelial cell line A549 expresses neither ACE2 or TMPRSS2, perhaps explaining why this cell line does not support infection by SARS-CoV-2 [12]. ACE2 and TMPRSS2 mRNA expression was also low in 293T cells, while ACE2 expression was considerably higher in 293T_ACE2_ cells (6074 fold increase from A549 cells). In contrast, HaCaT skin keratinocytes have the highest native ACE2 expression (277 fold increase from A549) but there was no expression of TMPRSS2 (Table 1; Fig 1). The Caco2 colorectal adenocarcinoma cell line, used in several infection studies of SARS-CoV and SARS-CoV-2 [13], was previously reported to express no ACE2 but high levels of TMPRSS2, however, qPCR analysis has demonstrated a 16 and 122 fold higher mRNA expression of ACE2 and TMPRSS2 respectively compared to A549 cells. Similarly, ACE2 and TMPRSS2 expression was 37 and 714 times higher respectively in the urinary bladder epithelial cell line, RT4 [14], compared to A549 cells (Fig 1). RT4 cells also express ADAM17, a metalloprotease known to be involved in the processing of ACE2, and CD9, an adaptor protein that controls ADAM17 trafficking and activity [15] (Table 1). Finally, RT4 cells also express rhomboid-like 2 (RHBDL2), a protease known to associate with ADAM17 [16], making this cell line a suitable choice for the study of viral attachment.

**Figure 1.**
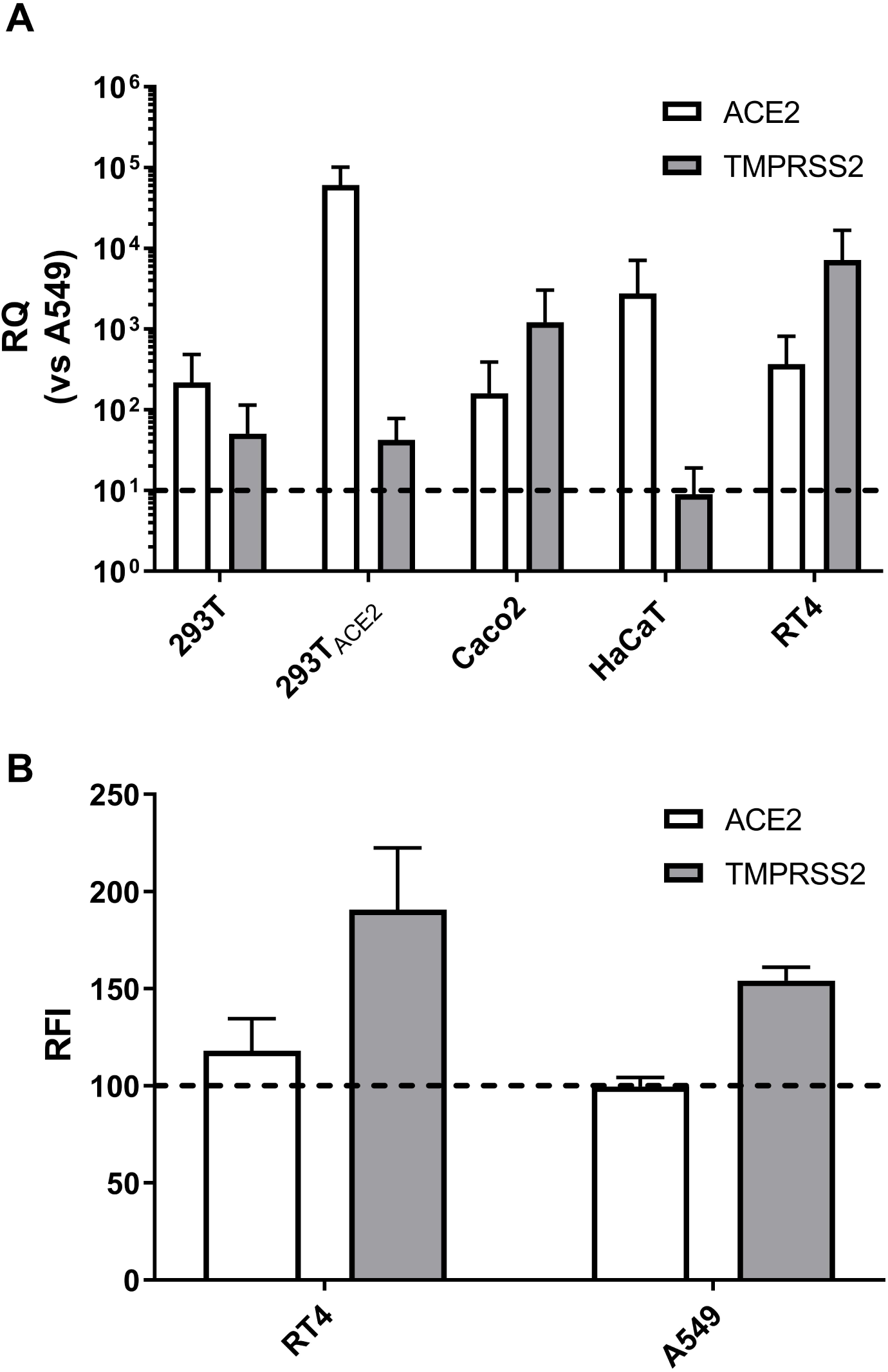
Expression of ACE2 on the surface of RT4 but not A549 cells. RT-qPCR and flow cytometry were used to compare the relative expression of ACE2 and TMPRSS2 across several human cell lines. To investigate gene expression levels, RNA was extracted, converted to cDNA and RT-qPCR was performed by SYBR Green assay. To investigate protein expression levels, RT4 and A549 cells were stained with goat anti-human ACE2 or rabbit anti-human TMPRSS2 antibodies and the appropriate fluorescent secondary antibodies. (A) Relative mRNA quantification of ACE2 (white) and TMPRSS2 (grey) in various cell lines compared to A549 cells (set at 10^1^), n≥2 (B) Relative fluorescence intensity of ACE2 and TMPRSS2 in A549 and RT4 cells. Relative fluorescence intensity was calculated as a percentage of negative control fluorescence, n≥2.

### Expression of ACE2 and TMPRSS2 at the surface of RT4 cells

We used flow cytometry to compare the expression of ACE2 and TMPRSS2 on RT4 and A549 cells. Despite high levels of mRNA within RT4 cells we observed only low levels of ACE2 at the surface of RT4 cells (Fig. 1), however, we cannot rule out the potential of a rapidly cycling population of ACE2 at the cell surface. TMPRSS2 expression was high in RT4 cells with little to no expression of either protein observed in A549 cells (Fig 1).

### Interaction of SARS-CoV-2 spike proteins with RT4 and A549 cells

To detect spike protein interactions, we used recombinant S1 tagged with mouse Fc and RBD or intact S1S2, both His6 tagged. Following published interaction studies for S1 [17], assays were initially performed at 21°C, using fluorescently labelled secondary antibodies to stain cells for flow cytometry. Only a very low level of S1 interacting with RT4 cells was detected, while S1 interaction with A549 cells was undetectable (S1 Fig). In contrast, S1S2 protein interacted strongly with subsets of both RT4 and A549 cells, however, a higher percentage of RT4 cells were positive compared to A549 cells (Fig 2A). Association with RT4 cells was detectable from 100nM S1S2 (Fig 2B) but was not saturated at 330nM, the highest concentration that could be used due to limited availability of the S1S2 protein.

**Figure 2.**
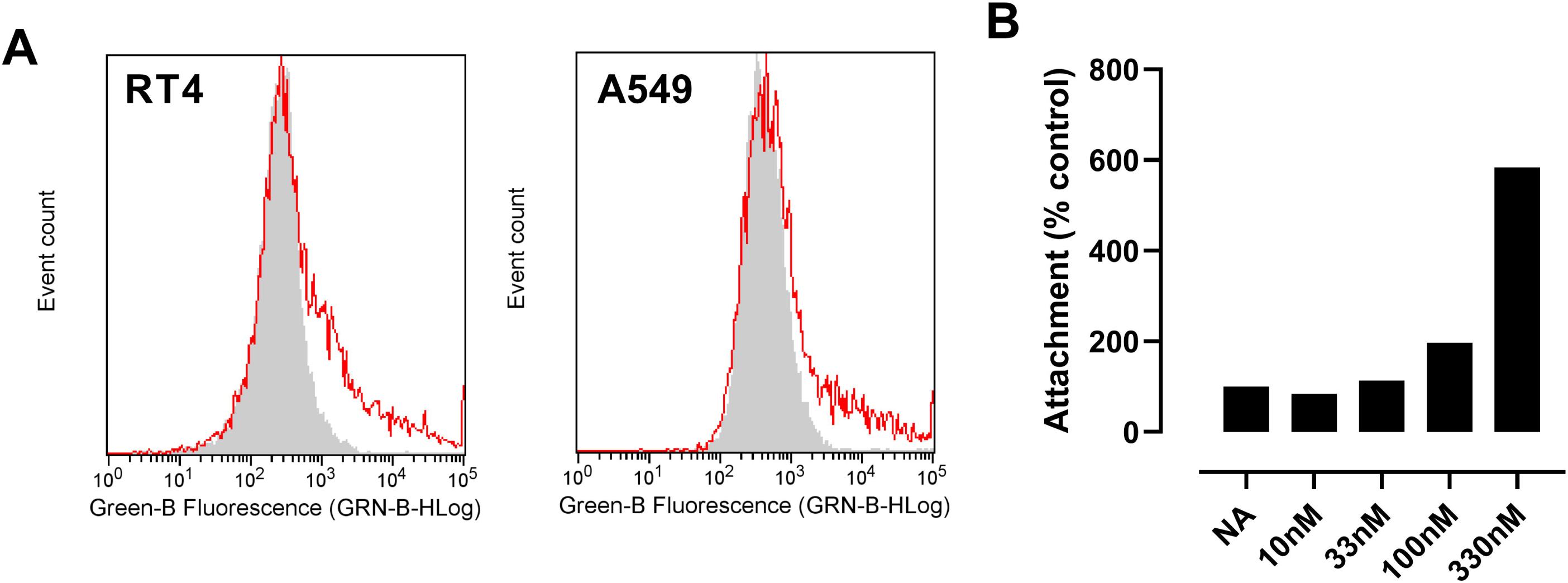
Recombinant S1S2 SARS-CoV-2 spike protein attaches to RT4 and A549 cells at 21°C. RT4 cells were incubated with His6-tagged S1S2 protein for 30 min at 21°C and then with anti-His6 secondary antibody labelled with Dylight 488. Cell associated fluorescence was measured by flow cytometry. (A) Representative attachment of 330nM S1S2 (red line) compared to secondary-only control (grey), n=2. (B) Attachment to RT4 cells measured as the number of cells more positive than the secondary antibody alone expressed as a percentage of the secondary-only controls (NA) for S1S2 from a single experiment conducted in duplicate.

To determine if the levels of detectable S1S2 interacting with RT4 cells were being affected by internalisation of the tagged protein, we performed assays at both 4°C, which should largely inhibit internalisation, and at 37°C, which should be permissive for internalisation. Surprisingly, interaction of S1S2 at 37°C was much stronger than at 4°C or 21°C (Fig 3), with all cells stained. In contrast, S1 or RBD protein association was undetectable at 4°C and only slightly elevated at 37°C (Fig 3). Interaction with A549 cells at 37°C was much lower than to RT4 cells (Fig 4A) at all concentrations tested although the cytometry histogram indicated that all cells had some S1S2 attached. This suggests that interaction at 37°C may be at least partly dependent on ACE2 and/or TMPRSS2 expression. S1S2 must be internalised only slowly, if at all, over the time course of the assay at 37°C. However, the temperature dependency suggests that S1S2 might undergo a conformational change, perhaps as a result of proteolytic processing at the cell surface. We were unable to determine the affinity of the interaction due to limited availability of recombinant S1S2, but association was still increasing even at 330nM (Fig 4C), suggesting the limiting step is a relatively low affinity interaction. This is in contrast to published reports of the affinity of S1 for 293T cells overexpressing human ACE2 (∼10nM) [17]. The polybasic site between the S1 and S2 domains (^681^PRRARSV^687^) is a putative furin cleavage site and has been shown to be essential for infection of human lung cells [2]. A mutant S1S2 (mS1S2) protein lacking the putative furin cleavage site demonstrated reduced interaction with RT4 cells compared to wtS1S2, similar to the S1 RBD alone (Fig4B, C). This might suggest the importance of proteolytic processing of the spike protein during viral association with the cell, although we cannot rule out the importance of a pair of stabilising modifications 300 amino acids C-terminal of the polybasic cleavage site within the mS1S2 [10] or potential differences in trimerization of the two proteins.

**Figure 3.**
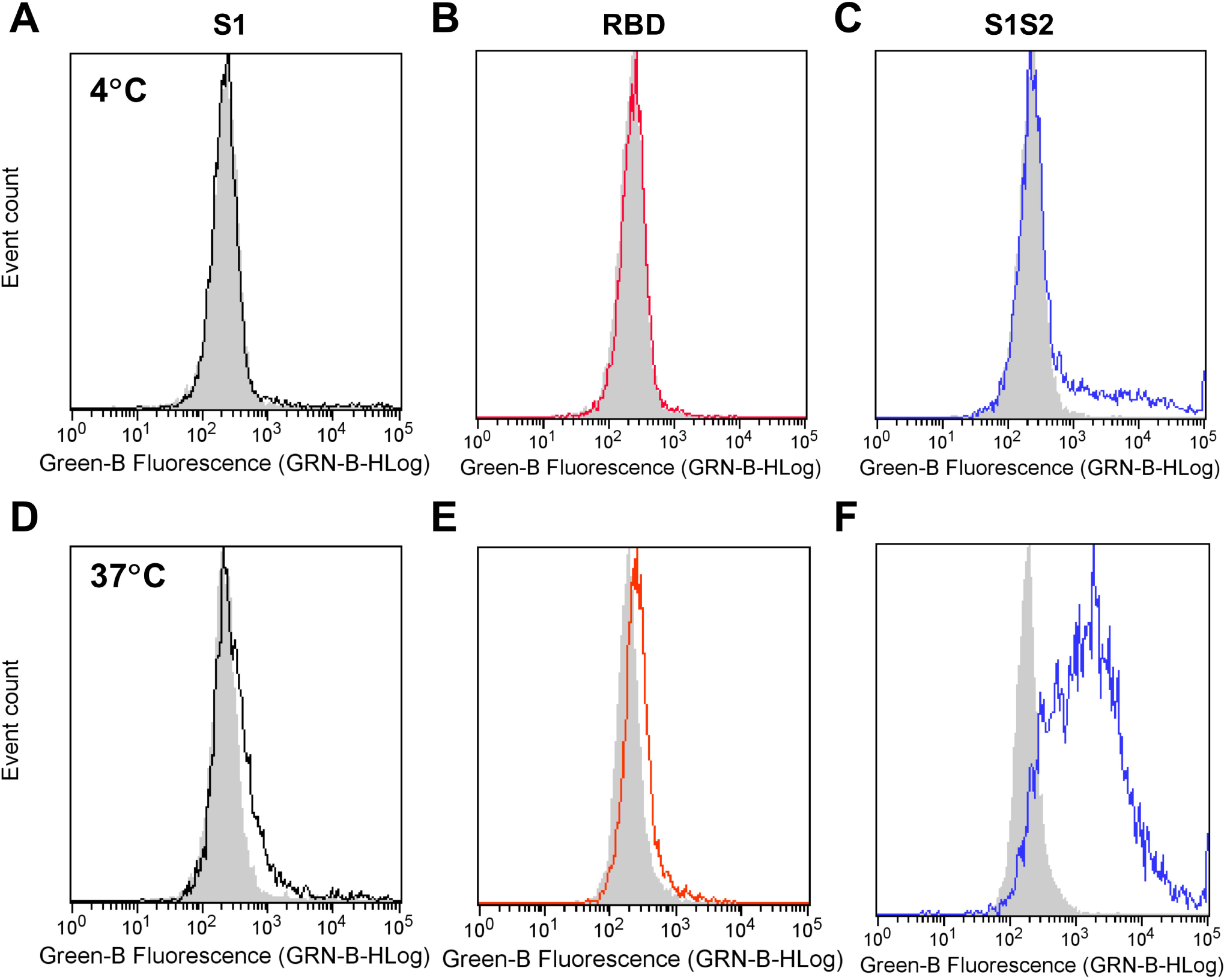
S1S2 attachment to RT4 cells dramatically increases at 37°C. RT4 cells were incubated with 330nM S1-Fc (A, D), 10µM RBD (B, E) or 330nM S1S2-His6 protein (C, F) for 60 mins at either 4°C (A-C) or 37°C (D-F), before staining with anti-mouse Ig labelled with FITC or anti-His6 secondary antibody labelled with Dylight 488 for 30 min at 21°C. Cell-associated fluorescence was measured using flow cytometry. Grey shows secondary-only control.

**Figure 4.**
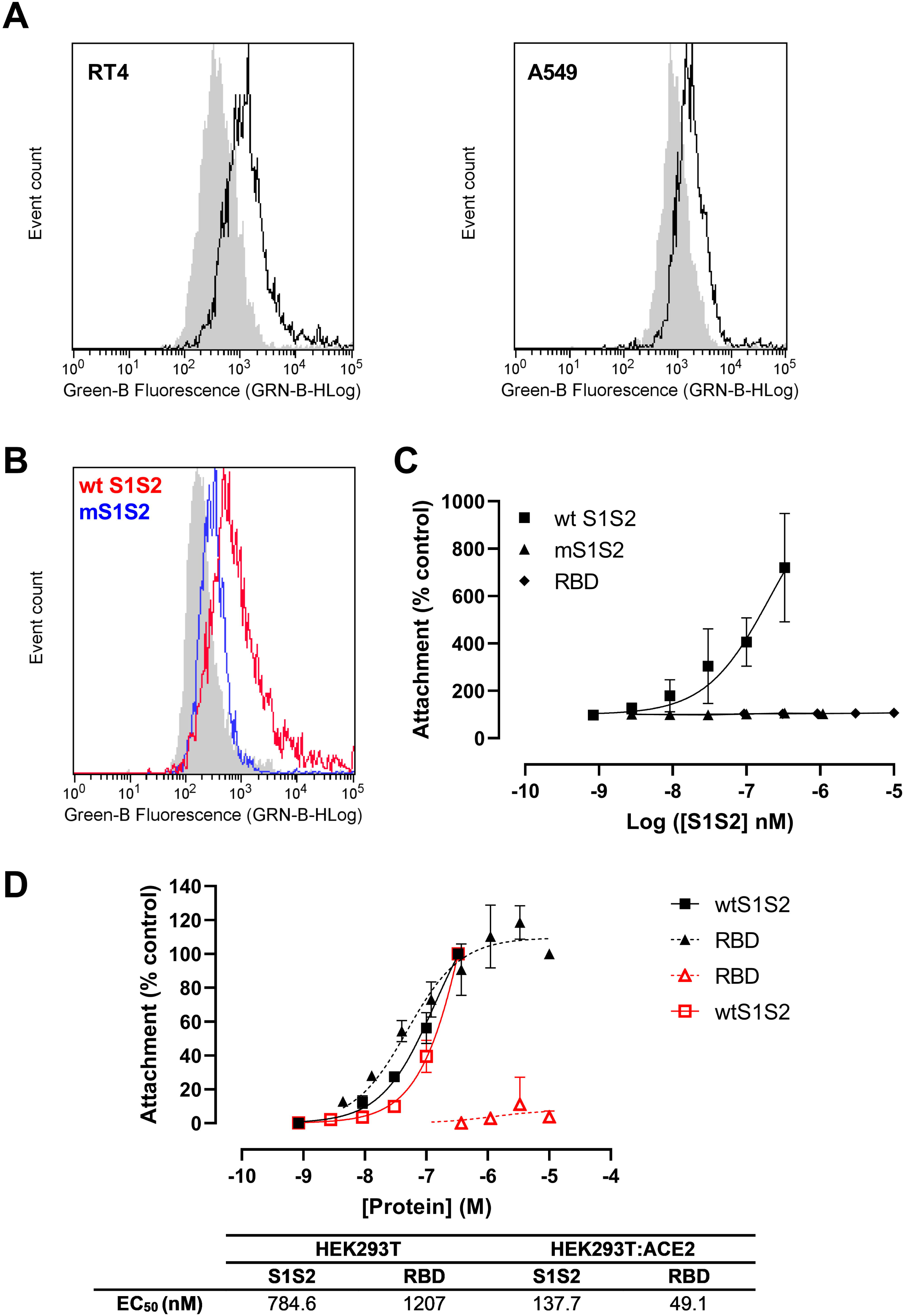
ACE2 increases S1S2 attachment to cells but is not required. Cells were incubated with wt S1S2 or RBD for 60 mins at 37°C, before staining with anti-His6 secondary antibody labelled with Dylight 488. (A) Representative histograms of 100nM S1S2 attachment to RT4 and A549 cells (black lines), compared to secondary antibody alone (grey). (B) Representative histogram of 100nM wild type S1S2 (wt S1S2) or mutant S1S2 (mS1S2) attachment to RT4 cells. (C) Dose-response curve for wt S1S2 (squares), mS1S2 (triangles) or receptor binding domain (RBD; diamonds) to RT4 cells. (D) Attachment of wt S1S2 (squares) or RBD (triangles) to 293T (red) or 293T_ACE2_ cells (black), n=3.

### S1S2 can interact with human cells independent of ACE2

As we observed some S1S2 interaction with A549 cells which lack ACE2, we sought to further investigate the role of spike protein interaction with ACE2 by assessing wtS1S2 and RBD attachment to 293T or 293T_ACE2_ cells (Fig 4D). wtS1S2 and RBD interacted with 293T cells overexpressing ACE2 with similar affinities (EC_50_ = 137.7nM and 49.1nM, respectively). In contrast, wtS1S2 associated with 293T cells expressing little ACE2, albeit at 5x lower affinity (EC_50_ = 784.6nM), while the affinity of the interaction of RBD with these cells was almost undetectable. This suggests that the initial interaction of S1S2 with cells is independent of ACE2, perhaps via an alternative receptor, whereas interaction of RBD alone requires high levels of cell surface ACE2.

### Unfractionated heparin inhibits S1S2 binding to RT4 cells

Having developed a novel assay that mimics some aspects of SARS-CoV-2 infection, we used it to test potential inhibitors. Although an anti-ACE2 antibody has previously been reported in one publication to block viral binding to host cells [1], preincubating RT4 cells with the same antibody caused a significant increase in S1S2 interaction (54.25% above untreated cells, p=0.0025) (Fig. 5A). Antibody-mediated cross-linking of ACE2 at the cell surface may allow rapid attachment of the spike protein due to receptor clustering. Heparin has also been reported to bind directly to S1 and to interfere with SARS-CoV-2 infection [7] and so we tested the effects of pre-incubating RT4 cells with heparin on the S1S2 attachment at 37°C. Unfractionated heparin (UFH) at 10U/ml inhibited 80% of 330nM S1S2 interaction with the cells (Fig 5B) and was significantly reduced compared to untreated controls (Fig 5C). Using 100nM S1S2, the inhibition by UFH was complete with an IC_50_ of 0.033U/ml (95% confidence interval 0.016-0.07) (Fig 6). This is far below the target prophylactic and therapeutic concentrations of UFH in serum, 0.1-0.4U/ml and 0.3-0.7U/ml, respectively [18,19]. Furthermore, the effect of UFH was also not caused by blockade of the His6 tag-antibody interaction (S2 Fig A), which was not affected in ELISA by concentrations of UFH of 500U/ml. In contrast, two low molecular weight heparins (LMWH), dalteparin and enoxaparin, were only partial inhibitors, and were less potent than UFH (IC_50_ values of 0.558 and 0.072U/ml, respectively). Typical prophylactic and therapeutic serum concentrations of LMWH are 0.2-0.5IU/ml and 0.5-1.2IU/ml [20], respectively, suggesting that dalteparin may be used below the effective dose required for inhibition of viral infection if used prophylactically. The synthetic pentasaccharide heparinoid, fondaparinux, had no effect on S1S2 interaction at concentrations up to 0.1mg/ml, although it has a therapeutic concentration of <2ug/ml [21] (Fig 6).

**Figure 5.**
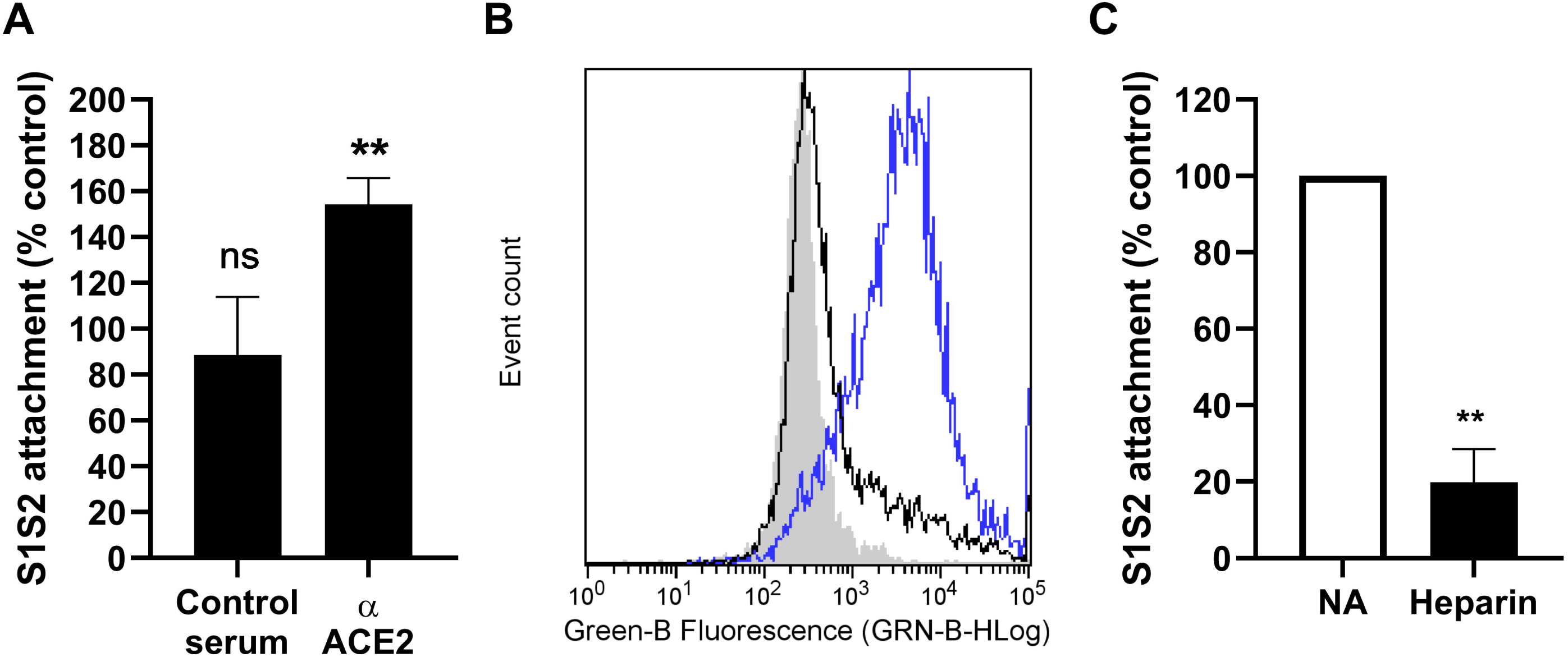
Unfractionated heparin inhibits S1S2 attachment to RT4 cells but anti-ACE2 antibody increases binding. RT4 cells were pre-treated with 15µg/ml anti-ACE-2 antibody (A) or 10U/ml unfractionated heparin (B-C) for 30 min at 37°C before the addition of 330nM S1S2. After a further 60 min at 37°C, cells were washed and fluorescent secondary anti-His6 added for a further 30 min at 21°C. Cell-associated fluorescence was measured by flow cytometry. (A) Effect of anti-ACE2 antibodies on 100nM S1S2 attachment, as a percentage of the S1S2 attachment to untreated control cells. Anti-goat IgG was used as a control. Significance to NA, ** p<0.01, one sample t-test, mean ± SEM, n=4. (B) Representative histogram with S1S2 attachment (blue line), S1S2 attachment after heparin treatment (black line) and secondary antibody only (grey). (C) Effect of 10U/ml heparin pre-treatment on 330nM S1S2 attachment, as a percentage of the S1S2 attachment to untreated (NA) control cells. Significance to NA, ** p<0.01, one sample t test, n=3.

**Figure 6.**
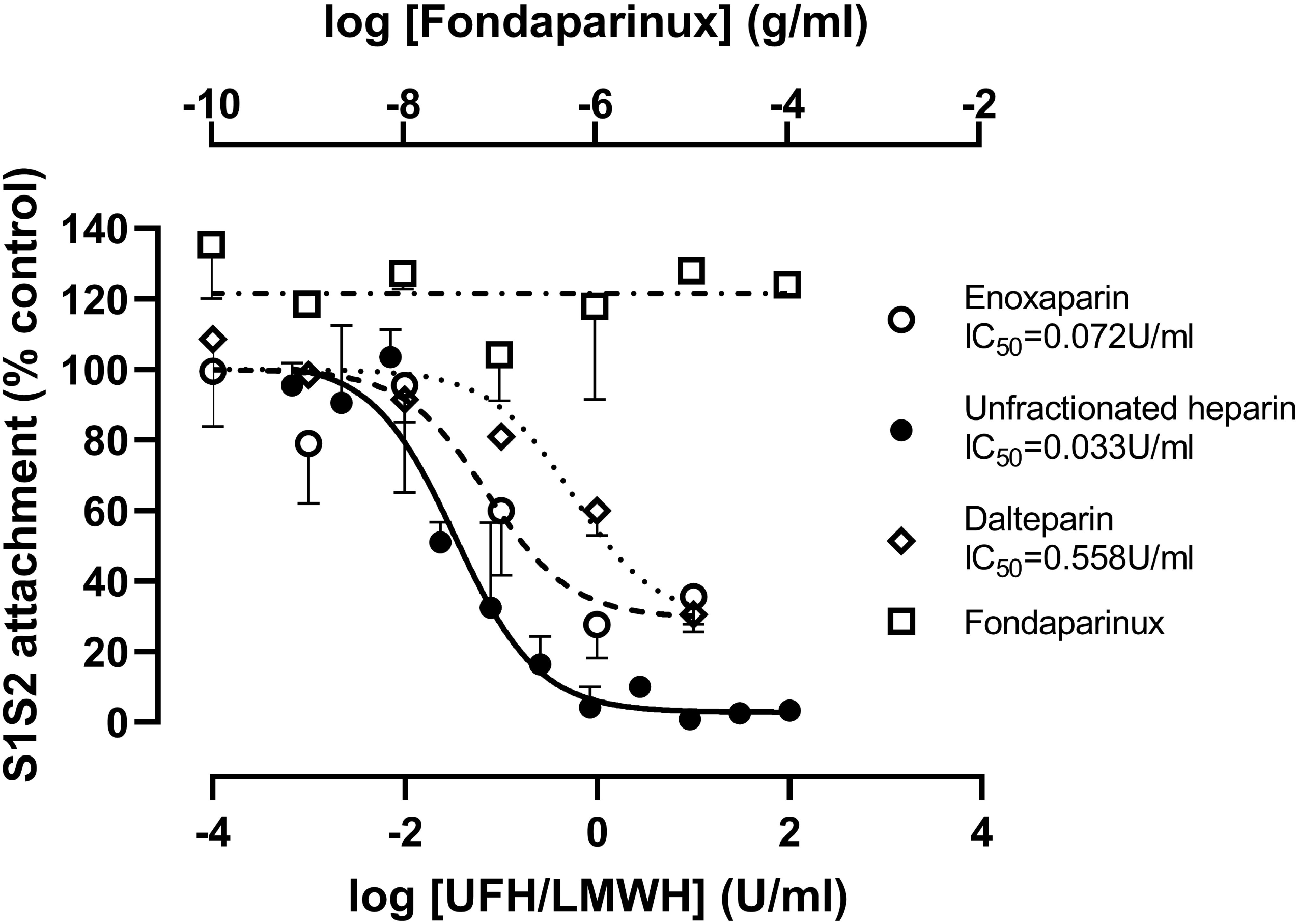
Concentration dependent inhibition of S1S2 attachment by unfractionated heparin and low molecular weight heparins, dalteparin and enoxaparin, but not synthetic pentasaccharide, fondaparinux. RT4 cells were pre-incubated with the stated concentrations of unfractionated heparin, enoxaparin, dalteparin and fondaparinux for 30 min at 37°C, then with 100nM S1S2 for a further 60 min at 37°C. After a further 60 min at 37°C, cells were washed and fluorescent secondary anti-His6 added for a further 30 min at 21°C. Cell-associated fluorescence was measured by flow cytometry and are shown as a percentage of the S1S2 attachment to untreated control cells. Data are the means ± SD of 2-3 experiments performed in duplicate.

### S1S2 that lacks the furin cleavage site has a lower affinity for heparin than wild type S1S2

The polybasic site furin cleavage site in S1S2 might also be a heparin binding site [22] and so we investigated if altered heparin binding by the mS1S2 might be linked to the lower binding to RT4 cells. mS1S2 and RBD had a significantly lower affinity for both UFH (mS1S2, EC_50_ = 217.8nM; RBD, EC_50_ = 818.4nM) and LMWH (mS1S2, EC_50_ = 162.2nM; RBD, EC_50_ = 288.3nM) relative to wild type (EC_50_ = 6.8nM and 9.3nM, respectively) (Fig 7). It is likely that wtS1S2 contains multiple binding sites for heparin that confer high avidity, and the polybasic furin cleavage site might be one of the most important of these sites. Additional sites may be present in S1 and RBD but these are of relatively low affinity. Interestingly, all the spike proteins bound UFH and dalteparin with similar affinities and so binding to immobilised heparins does not obviously explain the difference between UFH and LMWH in the inhibition of cell binding.

**Figure 7.**
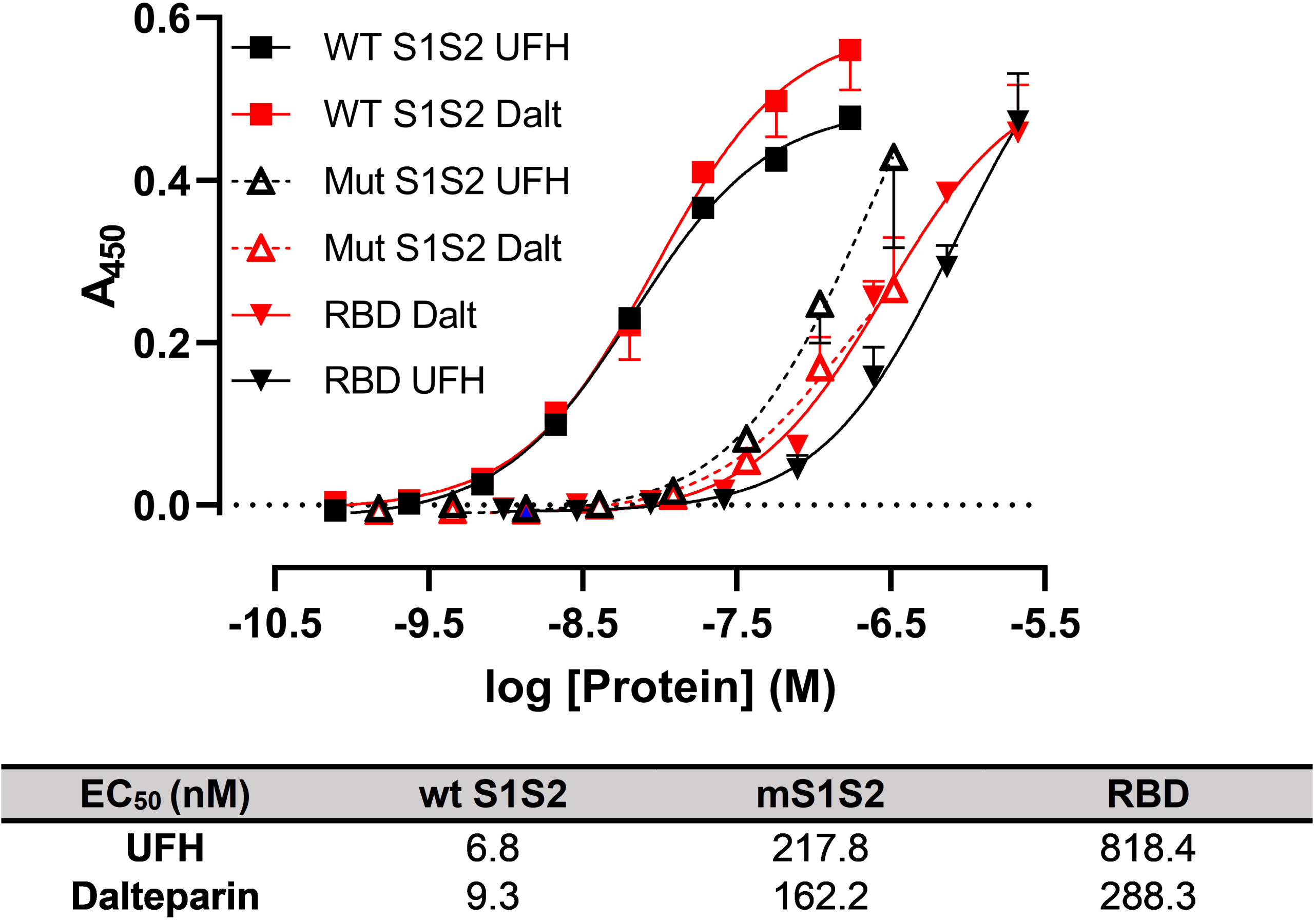
Spike protein binding to heparin requires a polybasic furin cleavage site. Unfractionated heparin (UFH) and low molecular weight heparin, dalteparin (Dalt) were immobilised on 96 well plates and used to detect the binding of His6-tagged wild type (WT S1S2), mutant S1S2 lacking the furin cleavage site (mS1S2) data and the receptor binding domain of S1 (RBD) using a biotinylated anti-His6 antibody and streptavidin-HRP. Data shown are the means ± SD from 2-3 separate experiments. Tables provide the EC_50_ for each protein.

## Discussion

We have demonstrated that intact recombinant wtS1S2 spike protein but not the S1 domain from SARS-CoV-2 can interact strongly with a human cell line that expresses ACE2 and TMPRSS2. Furthermore, S1S2 was able to associate with human cells independently of ACE2 expression, while the RBD alone could not. Using this assay, we have further shown that UFH and two low molecular weight heparins (LMWH) in use clinically can inhibit S1S2 binding. The same activity profile for UFH and one LMWH (enoxaparin) has been demonstrated in SARS-CoV-2 S1 RBD binding affinity [7]. These authors also showed that heparin could interact with recombinant S1 RBD and cause conformational changes, leading to the suggestion that SARS-CoV-2 might interact with host heparan sulphates through the RBD during infection.

Our data significantly extends this observation, suggesting the presence of multiple heparin binding sites in the intact spike protein, with one at the furin cleavage site and others in the S2 domain and within the RBD. As previously noted, the presence of multiple polybasic sites within the spike protein will result in a higher avidity of binding to heparin and heparan sulphates [9]. Interestingly, all the spike proteins tested had similar affinities for UFH and LMWH and so the differing abilities of these forms of heparin to inhibit S1S2 interactions with cells is not simply due to divergent affinities. However, we cannot rule out the possibility of differing glycosylation states affecting attachment of the recombinant proteins as their expression conditions varied.

Whilst previous studies have used African green monkey cells or human cells overexpressing relevant SARS-CoV-2 receptors, we have focused on native human cell lines. With reports of increased urinary frequency as an overlapping symptom of COVID-19 [23] and high expression of ACE2 and TMPRSS2 we have identified urinary bladder epithelial RT4 cells as a relevant model of infection for SARS-CoV-2. Whilst we demonstrate that ACE2 is involved, the intact spike protein was still able to interact with cells in the absence of ACE2 suggesting further secondary receptors. The ability of the SARS-CoV-2 spike protein to attach to various glycosaminoglycans provides a host of candidate proteins at the cell surface. Studies have also suggested the importance of differing glycan sulphation states in different tissues as an explanation for viral tropism. Recently, SARS-CoV-2 spike protein S1 has been shown to bind heparan sulphates with varying degrees of sulphation with differing affinities; chain length and 6-O-sulphation were particularly important [8].

UFH, and to a lesser extent, LMWH, are potent inhibitors of S1S2 interaction with human cells. LMWH are smaller (<8kDa) than UFH, which is a mix of polysaccharide chain lengths from ∼5-40kDa, and have more predictable pharmacokinetics [24]. LMWH are already commonly used both prophylactically and therapeutically in COVID-19 patients and have been reported to improve patient outcome [25]. Our work and the work of Mycroft-West *et al* suggests that thought be given to the earlier use of heparin when viral infection is still an important driver of disease severity. The use of UFH rather than LMWH should also be considered, although we note that administration and the safety profile of UFH might preclude this in some cases [26]. Fondaparinux contains only the antithrombin III interaction site and cannot form a ternary complex with thrombin, unlike UFH [21]. This suggests that efficient inhibition of S1S2 binding may require an interaction between heparin and two different sites on the spike protein. For example, SARS-CoV-2 spike protein optimally binds hexa- and octasaccharides composed of IdoA2S-GlcNS6S, a motif abundantly present within heparin but not heparan sulphates [9]. The LMWH may contain sufficient long polymer chains to make this dual interaction but with lower efficacy than UFH. Finally, heparin could also be inhibiting host proteases, like Factor Xa, necessary to process the spike protein [27]. However, our data demonstrated no activity of fondaparinux and lower efficacy of dalteparin despite the inhibitory nature of the two compounds on Factor Xa.

In conclusion, we have developed a simple flow cytometric assay for measuring SARS-CoV-2 spike protein interaction with human cells, demonstrating both S1S2 interaction with cells independent of ACE2 and potent inhibition of S1S2 association by UFH and LMWH. Our new assay could be a rapid initial screen for novel inhibitors of coronavirus infection.

## Acknowledgements

The authors would like to thank Dr Stephane Mesnage for flow cytometry, Prof. Florian Krammer for the kind gift of the mS1S2 and RBD plasmids, Dr. Thushan de Silva and Prof. Paul Bieniasz for the kind gift of the 293T_ACE2_ cells and the staff of the Molecular Biology and Biotechnology Department for access to laboratory space.

## Supporting information

**S1 Table. Primer Sequences**

**S1 Fig. S1-Fc attachment to RT4 and A549 cells is very low.** RT4 or A549 cells were incubated with 330nM mouse Fc-tagged S1 protein for 30 min at 21°C and then with anti-mouse Immunoglobulin secondary antibody labelled with FITC. Cell associated fluorescence was measured by flow cytometry. The histograms show S1 binding (black line) compared to secondary-only control (grey) and are representative of several separate experiments conducted in duplicate.

**S2 Fig. S1S2 attachment to ACE2 is not inhibited by unfractionated heparin.**

(A) Two His6-tagged proteins (human C5a and SARS-CoV-2 S1 RBD) were immobilised on the plate before visualisation using streptavidin-horseradish peroxidase and developed using TMB. The absorbance was measured at 450nm. Data are the results of single experiments performed in triplicate.

## References

1. Hoffmann M, Kleine-Weber H, Schroeder S, Krüger N, Herrler T, Erichsen S, et al. SARS-CoV-2 Cell Entry Depends on ACE2 and TMPRSS2 and Is Blocked by a Clinically Proven Protease Inhibitor. Cell. 2020;181: 271-280.e8. doi: 10.1016/j.cell.2020.02.052

2. Hoffmann M, Kleine-Weber H, Pöhlmann S. A Multibasic Cleavage Site in the Spike Protein of SARS-CoV-2 Is Essential for Infection of Human Lung Cells. Mol Cell. 2020;78: 779-784.e5. doi: 10.1016/j.molcel.2020.04.022

3. Hofmann H, Pöhlmann S. Cellular entry of the SARS coronavirus. Trends Microbiol. 2004;12: 466–472. doi: 10.1016/j.tim.2004.08.008

4. Li MY, Li L, Zhang Y, Wang XS. Expression of the SARS-CoV-2 cell receptor gene ACE2 in a wide variety of human tissues. Infect Dis Poverty. 2020;9: 45. doi: 10.1186/s40249-020-00662-x

5. O’Donnell CD, Shukla D. The importance of heparan sulfate in herpesvirus infection. Virol Sin. 2008;23: 383–393. doi: 10.1007/s12250-008-2992-1

6. Kim SY, Jin W, Sood A, Montgomery DW, Grant OC, Fuster MM, et al. Characterization of heparin and severe acute respiratory syndrome-related coronavirus 2 (SARS-CoV-2) spike glycoprotein binding interactions. Antiviral Res. 2020;181: 104873. doi: 10.1016/j.antiviral.2020.104873

7. Mycroft-West CJ, Su D, Pagani I, Rudd TR, Elli S, Guimond SE, et al. Heparin inhibits cellular invasion by SARS-CoV-2: structural dependence of the interaction of the surface protein (spike) S1 receptor binding domain with heparin. bioRxiv. 2020; 2020.04.28.066761. doi: 10.1101/2020.04.28.066761

8. Hao W, Ma B, Li Z, Wang X, Gao X, Li Y, et al. Binding of the SARS-CoV-2 Spike Protein to Glycans. bioRxiv. 2020; 2020.05.17.100537. doi: 10.1101/2020.05.17.100537

9. Liu L, Chopra P, Li X, Wolfert MA, Tompkins SM, Boons G-J. SARS-CoV-2 spike protein binds heparan sulfate in a length- and sequence-dependent manner. bioRxiv. 2020; 2020.05.10.087288. doi: 10.1101/2020.05.10.087288

10. Amanat F, Stadlbauer D, Strohmeier S, Nguyen THO, Chromikova V, McMahon M, et al. A serological assay to detect SARS-CoV-2 seroconversion in humans. Nat Med. 2020. doi: 10.1038/s41591-020-0913-5

11. Uhlen M, Zhang C, Lee S, Sjöstedt E, Fagerberg L, Bidkhori G, et al. A pathology atlas of the human cancer transcriptome. Science (80-). 2017;357. doi: 10.1126/science.aan2507

12. Matsuyama S, Nao N, Shirato K, Kawase M, Saito S, Takayama I, et al. Enhanced isolation of SARS-CoV-2 by TMPRSS2-expressing cells. Proc Natl Acad Sci U S A. 2020;117: 7001–7003. doi: 10.1073/pnas.2002589117

13. Kim JM, Chung YS, Jo HJ, Lee NJ, Kim MS, Woo SH, et al. Identification of coronavirus isolated from a patient in Korea with covid-19. Osong Public Heal Res Perspect. 2020;11: 3–7. doi: 10.24171/j.phrp.2020.11.1.02

14. O’Toole C, Perlmann P, Unsgaard B, Almgård LE, Johansson B, Moberger G, et al. Cellular immunity to human urinary bladder carcinoma. II. Effect of surgery and preoperative irradiation. Int J Cancer. 1972;10: 92–98. doi: 10.1002/ijc.2910100112

15. Gutiérrez-López MD, Gilsanz A, Yáñez-Mó M, Ovalle S, Lafuente EM, Domínguez C, et al. The sheddase activity of ADAM17/TACE is regulated by the tetraspanin CD9. Cell Mol Life Sci. 2011;68: 3275–3292. doi: 10.1007/s00018-011-0639-0

16. McIlwain DR, Lang PA, Maretzky T, Hamada K, Ohishi K, Maney SK, et al. iRhom2 regulation of TACE controls TNF-mediated protection against Listeria and responses to LPS. Science (80-). 2012;335: 229–232. doi: 10.1126/science.1214448

17. Tai W, Zhang X, He Y, Jiang S, Du L. Identification of SARS-CoV RBD-targeting monoclonal antibodies with cross-reactive or neutralizing activity against SARS-CoV-2. Antiviral Res. 2020;179: 104820. doi: 10.1016/j.antiviral.2020.104820

18. Hirsh J, Warkentin TE, Shaughnessy SG, Anand SS, Halperin JL, Raschke R, et al. Heparin and low-molecular-weight heparin: Mechanisms of action, pharmacokinetics, dosing, monitoring, efficacy, and safety. Chest. 2001;119: 64S–94S. doi: 10.1378/chest.119.1_suppl.64S

19. Lehman CM, Frank EL. Laboratory Monitoring of Heparin Therapy: Partial Thromboplastin Time or Anti-Xa Assay? Lab Med. 2009;40: 47–51. doi: 10.1309/lm9njgw2ziolphy6

20. Jaffer IH, Weitz JI. Antithrombotic Drugs. Hematology: Basic Principles and Practice. Elsevier Inc.; 2018. pp. 2168–2188. doi: 10.1016/B978-0-323-35762-3.00149-9

21. Bauer KA, Hawkins DW, Peters PC, Petitou M, Herbert JM, Van Boeckel AA, et al. Fondaparinux, a synthetic pentasaccharide: The first in a new class of antithrombotic agents - The selective factor Xa inhibitors. Cardiovascular Drug Reviews. 2002. pp. 37–52. doi: 10.1111/j.1527-3466.2002.tb00081.x

22. Muñoz EM, Linhardt RJ. Heparin-binding domains in vascular biology. Arteriosclerosis, Thrombosis, and Vascular Biology. NIH Public Access; 2004. pp. 1549–1557. doi: 10.1161/01.ATV.0000137189.22999.3f

23. Mumm J-N, Osterman A, Ruzicka M, Stihl C, Vilsmaier T, Munker D, et al. Urinary Frequency as a Possibly Overlooked Symptom in COVID-19 Patients: Does SARS-CoV-2 Cause Viral Cystitis? Eur Urol. 2020 [cited 8 Jul 2020]. doi: 10.1016/j.eururo.2020.05.013

24. Samama M, Bernard P, Bonnardot JP, Combe□Tamzali S, Lanson Y, Tissot E. Low molecular weight heparin compared with unfractionated heparin in prevention of postoperative thrombosis. Br J Surg. 1988;75: 128–131. doi: 10.1002/bjs.1800750213

25. Tang N, Bai H, Chen X, Gong J, Li D, Sun Z. Anticoagulant treatment is associated with decreased mortality in severe coronavirus disease 2019 patients with coagulopathy. J Thromb Haemost. 2020;18: 1094–1099. doi: 10.1111/jth.14817

26. Junqueira DR, Zorzela LM, Perini E. Unfractionated heparin versus low molecular weight heparins for avoiding heparin-induced thrombocytopenia in postoperative patients. Cochrane Database Syst Rev. 2017;2017. doi: 10.1002/14651858.CD007557.pub3

27. Belen-Apak FB, Sarialioglu F. The old but new: Can unfractioned heparin and low molecular weight heparins inhibit proteolytic activation and cellular internalization of SARS-CoV2 by inhibition of host cell proteases? Med Hypotheses. 2020;142. doi: 10.1016/j.mehy.2020.109743

