## Supplementary material for "ACE2-independent interaction of SARS-CoV-2 spike protein to human epithelial cells can be inhibited by unfractionated heparin": S1 Table

| **Primer name** | **Sequence (5’-3’)** |
| --- | --- |
| ACE2 F | GGGATCAGAGATCGGAAGAAGAAA |
| ACE2 R | AGGAGGTCTGAACATCATCAGTG |
| TMPRSS2 F | TAGTGTCCCCAGCCTACCTC |
| TMPRSS2 R | GCACCAAGGGCACTGTCTAT |
| GAPDH F | TGCACCACCAACTGCTTAGC |
| GAPDH R | GGCATGGACTGTGGTCATGAG |
