## Supplementary figures and images for "ACE2-independent interaction of SARS-CoV-2 spike protein to human epithelial cells can be inhibited by unfractionated heparin"

### S1 Fig

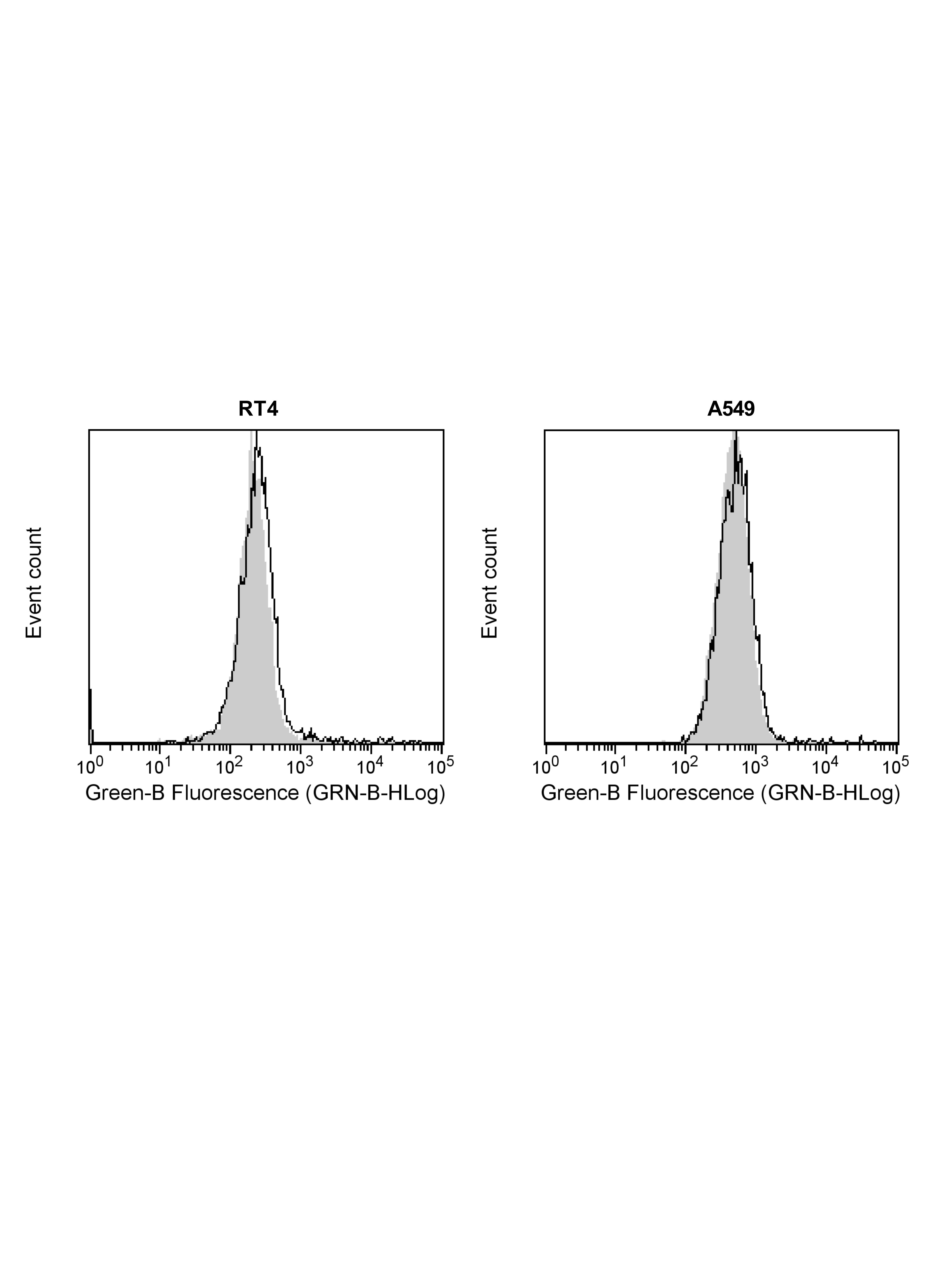

### S2 Fig

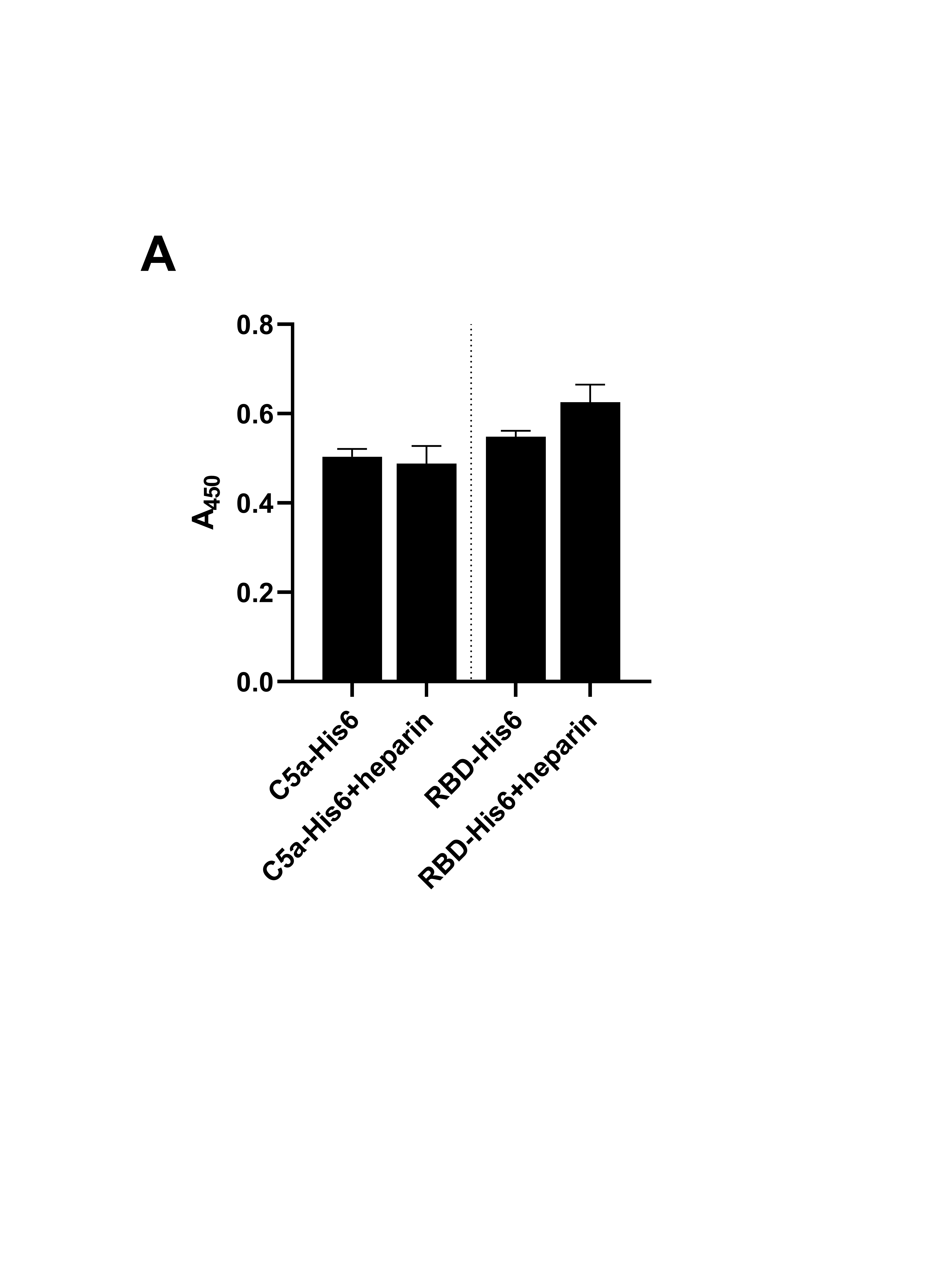
